# The crossover function of MutSγ is activated via Cdc7-dependent stabilization of Msh4

**DOI:** 10.1101/386458

**Authors:** Wei He, H.B.D. Prasada Rao, Shangming Tang, Nikhil Bhagwat, Dhananjaya S. Kulkarni, Maria A.W. Chang, Christie Hall, Lepakshi Singh, Xiangyu Chen, Nancy M. Hollingsworth, Petr Cejka, Neil Hunter

## Abstract

The MutSγ complex, Msh4-Msh5, binds DNA joint-molecule (JM) intermediates during homologous recombination to promote crossing over and accurate chromosome segregation at the first division of meiosis. MutSγ facilitates the formation and biased resolution of crossover-specific JM intermediates called double Holliday junctions. Here we show that these activities are governed by regulated proteasomal degradation. MutSγ is initially inactive for crossing over due to an N-terminal degron on Msh4 that renders it unstable. Activation of MutSγ requires the Dbf4-dependent kinase, Cdc7 (DDK), which directly phosphorylates and thereby neutralizes the Msh4 degron. Phosphorylated Msh4 is chromatin bound and requires DNA strand exchange and chromosome synapsis, implying that DDK specifically targets MutSγ that has already bound nascent JMs. Our study establishes regulated protein degradation as a fundamental mechanism underlying meiotic crossover control.

## INTRODUCTION

Crossing over is required for the accurate segregation of homologous chromosomes (homologs) at the first division of meiosis (Hunter, 2015; Watanabe, 2012). At metaphase I, the four chromatids of a bivalent are interconnected by cohesion between sister chromatids and at least one crossover between the homologs. These connections enable stable bipolar attachment of bivalents to the meiosis-I spindle, which is a prerequisite for accurate homolog disjunction at the ensuing anaphase. Crossing over is highly regulated to ensure that homolog pairs obtain the requisite connection despite a low average number of crossovers per nucleus (Jones, 1984). Also, crossovers between a single homolog pair inhibit one another such that multiple events tend to be widely and evenly spaced (Wang et al., 2015). Together, these crossover assurance and crossover interference processes dictate the range of crossover numbers per nucleus. In *C. elegans*, interference is effective along the entire lengths of all chromosomes, minimizing crossover numbers to one per chromosome (Hillers and Villeneuve, 2003). More typically, crossovers average around one per chromosome arm, but smaller chromosomes will often obtain only a single exchange. Suboptimal crossing-over, leading to missegregation and aneuploidy, and aberrant exchange between non-allelic homologies are leading causes of congenital disease in humans (Herbert et al., 2015; Hunter, 2015; Kim et al., 2016; Wang et al., 2017).

The regulatory processes that underlie crossover control remain poorly understood. Crossover sites are designated from a larger pool of recombination sites initiated by Spo11-catalyzed DNA double-strand breaks (DSBs)(Lam and Keeney, 2014). DSBs outnumber crossovers by ∼2-fold in budding yeast, ∼10-fold in mammals and up to 30-fold in some plants. At the cytological level, the differentiation of crossover sites manifests as the selective retention and accumulation of specific recombination factors. One such factor is MutSγ, a heterodimer of Msh4 and Msh5, two homologs of the DNA mismatch-recognition factor MutS (Manhart and Alani, 2016; Pochart et al., 1997; Snowden et al., 2004). Msh4 and Msh5 are members of the ZMM proteins (Zip1, Zip2, Zip3, Zip4, Msh4, Msh5, Mer3, and Spo16), a diverse set of factors that facilitate crossover-specific recombination events, and couple these events to chromosome synapsis (Borner et al., 2004; Fung et al., 2004; Hunter, 2015; Lynn et al., 2007; Shinohara et al., 2008). As seen in a variety of species, initial numbers of MutSγ immunostaining foci greatly outnumber final crossover numbers (De Muyt et al., 2014; de Vries et al., 1999; Edelmann et al., 1999; Higgins et al., 2008; Kneitz et al., 2000; Yokoo et al., 2012; Zhang et al., 2014). As prophase I progresses, MutSγ is lost from most recombination sites but retained at sites that mature into crossovers. This patterning process is dependent on the Zip3/RNF212/ZHP-3/HEI10 family of RING E3 ligases (Agarwal and Roeder, 2000; Bhalla et al., 2008; Chelysheva et al., 2012; De Muyt et al., 2014; Henderson and Keeney, 2004; Jantsch et al., 2004; Qiao et al., 2014; Rao et al., 2017; Reynolds et al., 2013; Wang et al., 2012; Yokoo et al., 2012; Zhang et al., 2018). Additional evidence implicates the SUMO-modification and ubiquitin-proteasome systems in meiotic crossover control (Ahuja et al., 2017; Rao et al., 2017) and suggests a model in which factors such as MutSγ are selectively stabilized at crossover sites by protecting them from proteolysis. Implicit in this model is the notion that MutSγ is intrinsically unstable.

Here, we show that regulated proteolysis plays a direct and essential role in meiotic crossing over. Msh4 is identified as an intrinsically unstable protein that is targeted for proteasomal degradation by an N-terminal degron thereby rendering MutSγ inactive for crossing over. Activation of MutSγ occurs by neutralizing the Msh4 degron via phosphorylation catalyzed by the conserved cell-cycle kinase, Cdc7-Dbf4 (DDK). Thus, a key meiotic pro-crossover factor is activated by attenuating its proteolysis.

## RESULTS

### The N-terminal Region of Msh4 is Phosphorylated

The ZMM proteins were surveyed for modifications detectable as electrophoretic-mobility shifts on Western blots. A prominent modified band was detected for Msh4 but not for its partner Msh5 (**Figure 1A**). Treatment of immunoprecipitated Msh4 with λ phosphatase indicated that the modified form is due to phosphorylation (**Figure 1B**). Relative to the unphosphorylated protein, phosphorylated Msh4 appeared with a ≥1 hr delay, its levels peaked at ∼22% of total protein, and then both species disappeared with the same timing (**Figure 1C**). To map sites of phosphorylation, Msh4 was immunoprecipitated, fast and slow migrating forms were resolved by electrophoresis, and then analyzed separately by tandem mass spectrometry (MS/MS; **Figure 1D**). Six phosphorylation sites were identified in the slower migrating form of Msh4, all mapping within the first 50 amino acids (S2, S4, S7, S41, T43 and S46; **Figure 1E; Supplemental Figure S1**). In the faster migrating form of Msh4, only phosphorylation at S41 was detected.

**Figure 1.**
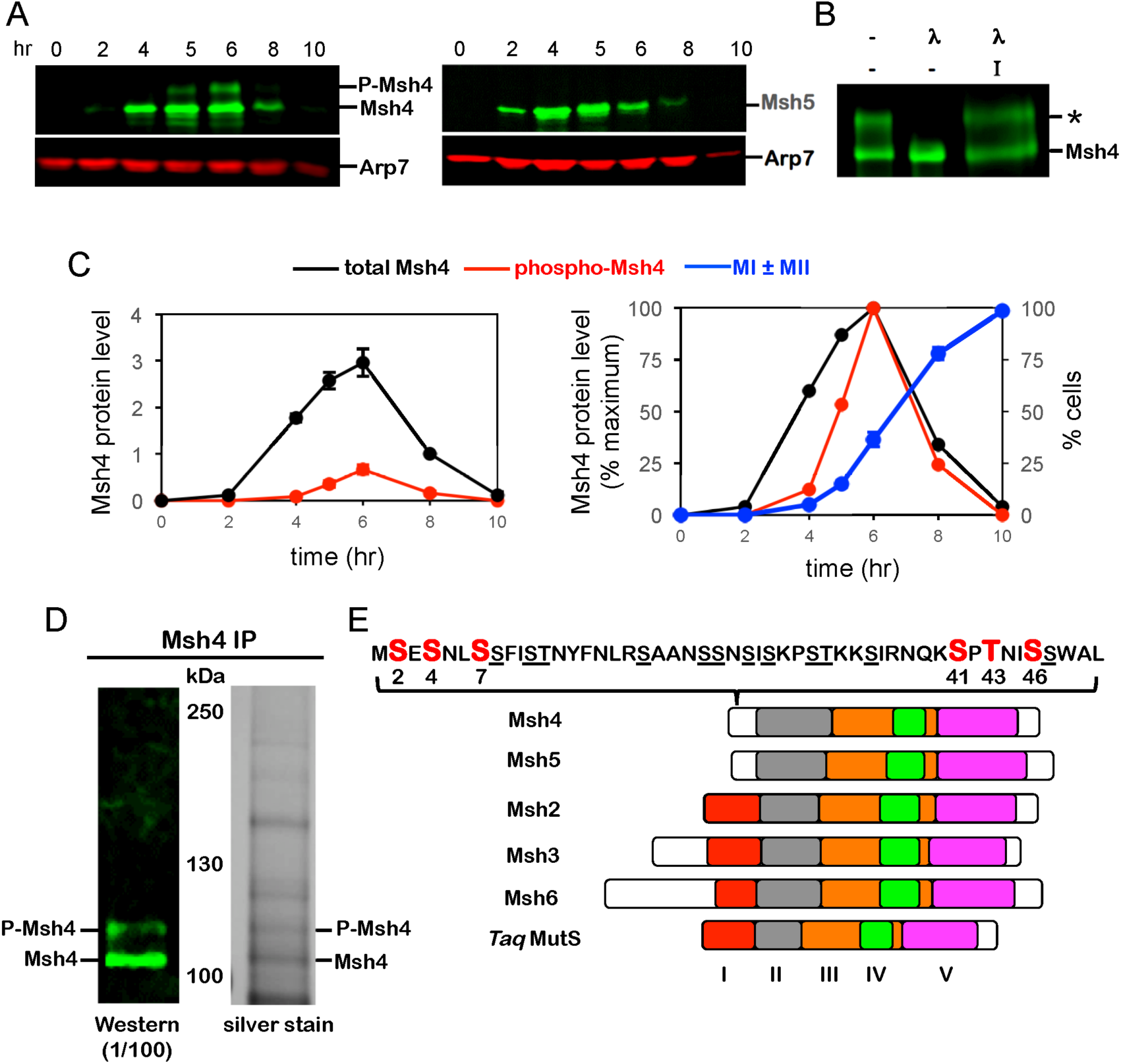
The N-terminal region of Msh4 is phosphorylated. (A) Western analysis of Msh4 (left) and Msh5 (right) throughout meiosis. Arp7 is used throughout as a loading control (Sourirajan and Lichten, 2008). (B) Lambda phosphatase treatment of immuno-precipitated Msh4. λ, phosphatase; I, phosphatase inhibitor. (C) Relative (left) and normalized (right) levels Msh4 phosphorylation. MI ± MII is the percentage of cells that have completed one or both meiotic divisions. Error bars show mean ± S.E. from four independent time courses. (D) Western blotting and silver-stained gel images of immunoprecipitated material used for LC-MS/MS analysis. The positions of the Msh4 bands that were excised and processed for LC-MS/MS are indicated. (E) Positions of phosphorylation sites (red) mapped by LC-MS/MS. Underlined residues highlight the high S/T content of the Msh4 N-terminal region. Diagrams show protein domains of eukaryotic nuclear MutS homologs relative to *Thermus aquaticus* MutS (adapted from (Nishant et al., 2010). Also see **Figure S1**.

Msh4 and Msh5 lack the N-terminal domain I, which is conserved in other MutS proteins (**Figure 1E**). Domain I encircles DNA together with MutS domain IV and is intimately involved in DNA binding and mismatch recognition (Yang et al., 2000). Absence of domain I from Msh4 and Msh5 is predicted to enlarge the DNA channel such that it can accommodate JM structures and slide on two duplexes (Rakshambikai et al., 2013; Snowden et al., 2004). The functions of the N-terminal regions of Msh4 and Msh5 are otherwise unknown.

### Phosphorylation is Essential for the Crossover Function of Msh4

To determine the functional relevance of Msh4 phosphorylation, we mutated the six identified phosphorylation sites to alanine to prevent phosphorylation, or to aspartic acid to mimic phosphorylation, creating respectively *msh4-6A* and *msh4-6D* alleles (**Figure 2**). Spore viability, indicative of successful meiotic chromosome segregation, was assessed by tetrad dissection and compared to wild-type and *msh4*Δ-null mutant strains (**Figure 2A,B**). Consistent with previous studies (Krishnaprasad et al., 2015; Nishant et al., 2010; Novak et al., 2001; Stahl et al., 2004), *msh4*Δ reduced spore viability to 34.7% and the pattern of spore death was indicative of chromosome missegregation at the first meiotic division, with a preponderance of tetrads containing two or zero viable spores (**Figure 2B**). The pattern of spore death in cells carrying the phosphorylation-defective *msh4-6A* allele was similar to that of the *msh4*Δ null, with an overall viability of 46.7% (*P*<0.01 compared to wild type, 𝒳^2^ test; **Table S1**). By contrast, the phospho-mimetic *msh4-6D* allele supported wild-type levels of spore viability (96.3% and 95.7%, respectively, *P*=0.42).

**Figure 2.**
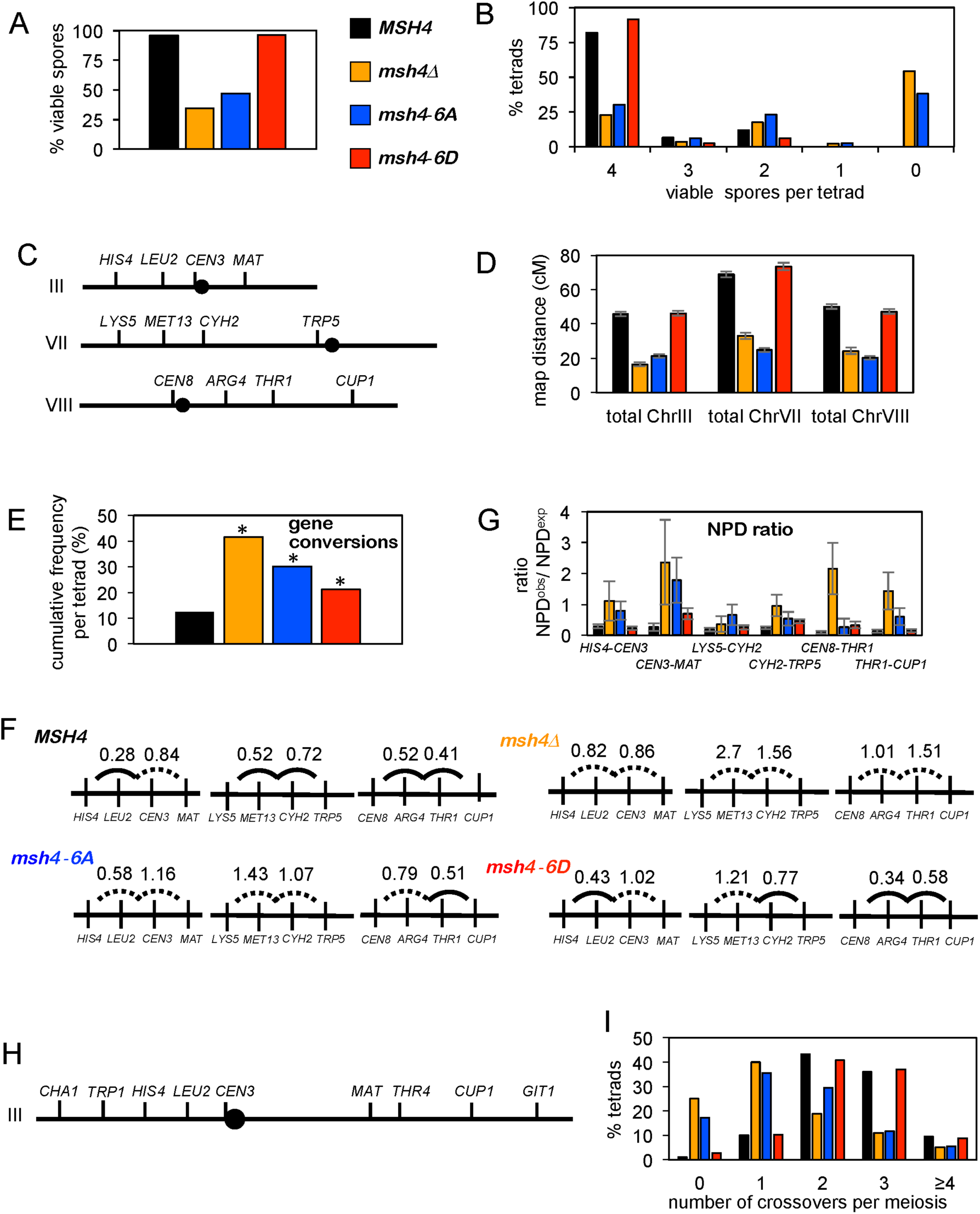
Phosphorylation is essential for the crossover function of Msh4. (A) Spore viabilities of indicated strains (see **Table S3**). (B) Distributions of tetrads with 4, 3, 2, 1 and 0 viable spores. (C) Marker configurations in strains used to analyze recombination. *CEN3* is marked with the *ADE2* gene and *CEN8* is marked with *URA3* (Oh et al., 2007). (D) Cumulative map distances (± S.E.) for intervals on chromosomes III, VII and VIII. (E) Cumulative frequencies of tetrads with gene conversions (non 2:2 segregations) for markers shown in (E). Asterisks indicate *P*<0.01 relative to wild type (*z*-test). (F) Interference analysis for adjacent intervals (Lao et al., 2013; Malkova et al., 2004). Solid arcs between intervals indicate significant positive interference; failure to detect significant positive interference is indicated by dashed arcs (see **Table S4**). (G) Interference analysis within individual intervals expressed as NPD ratios (https://elizabethhousworth.com/StahlLabOnlineTools/). Error bars show S.E. Asterisks indicate significant positive interference (see **Table S6**). (H) Marker configuration in strains used to analyze crossover assurance for chromosome III. *CEN3* is heterozgously marked with the *LYS2 and URA3* (see **Table S1).** (I) Distributions of crossover classes for chromosome III. See also **Figures S2** and **S3**.

In the absence of Msh4, chromosome missegregation and the ensuing spore death are caused by defective crossing over (Krishnaprasad et al., 2015; Nishant et al., 2010; Novak et al., 2001; Stahl et al., 2004). To assess whether phosphorylation is required for the crossover function of Msh4, we measured genetic map distances in a background carrying markers on three different chromosomes (III, VII and VIII; **Figure 2C**). Cumulative map distances showed that *msh4*Δ reduced crossing by 2.1 to 2.7-fold (**Figure 2D** and **Supplemental Figure S2** and **Table S2**). Similar reductions (2.1 to 2.8-fold) were seen for the *msh4-6A* phosphorylation-defective strain. Thus, phosphorylation is essential for the crossover function of Msh4. For chromosomes VII and VIII, the *msh4-6A* mutation caused slightly larger reductions in crossing over than the *msh4*Δ null. Possibly, phosphorylation-defective Msh4-6A protein is still capable of binding recombination intermediates thereby impeding processing via alternative crossover pathways mediated by the structure-selective nucleases (De Muyt et al., 2012; Zakharyevich et al., 2012). Consistent with the high spore viability of the phospho-mimetic *msh4-6D* strain, cumulative map distances for this strain were indistinguishable from those of wild type.

Previous analysis showed that non-crossover gene conversions are increased in the absence of ZMMs, including Msh5 (and by extension Msh4), due to the continued formation of DSBs when homolog engagement is defective (Hollingsworth et al., 1995; Novak et al., 2001; Thacker et al., 2014). This phenotype was reflected in tetrad data from the *msh4*Δ null strain, which showed a 3.4-fold increase in the cumulative gene conversion frequency for the 12 markers in this background (**Figure 2E**; also see **Supplemental Figure S2**). Elevated gene conversion was also seen for *msh4-6A,* which showed a 2.5-fold increase in gene conversions relative to wild type. Unexpectedly, a 1.7-fold increase in gene conversion was observed for the *msh4-6D* strain, the first indication that this phospho-mimetic allele does not possess fully wild-type function.

Crossovers promoted by MutSγ are patterned by interference (Krishnaprasad et al., 2015; Novak et al., 2001; Stahl et al., 2004). One readout of crossover interference is that tetrads with crossovers in a given “test” interval have significantly lower frequencies of crossovers in the neighboring intervals when compared to tetrads that lack crossovers in the test interval (Malkova et al., 2004)(**Figure 2F**). This difference can be expressed as the ratio of map distances for the neighboring interval in the tetrad subsets with or without a crossover in the test interval. Positive crossover interference is indicated by a ratio that is significantly less than one. To address whether Msh4 phosphorylation promotes crossovers with an interference distribution, this analysis was performed for all interval pairs (**Figure 2F**; also see **Supplemental Figure S3**).

In wild-type tetrads, significant positive crossover interference was detected for all interval pairs except *LEU2–CEN3–MAT* on chromosome 3 (**Figure 2F**). Consistent with previous studies (Novak et al., 2001; Stahl et al., 2004), residual crossovers in the *msh4*Δ strain did not show significant positive interference in any interval pair. In fact, significant negative interference – a higher incidence of double crossovers than expected – was detected for one interval pair on chromosome VII (*LYS5-MET13-CYH2*) in *msh4*Δ tetrads. Analogous results were obtained for the *msh4-6A* strain indicating that Msh4 phosphorylation does indeed promote the formation of crossovers that are subject to interference. However, significant positive interference was still detected in one interval pair for *msh4-6A* (*ARG4-THR1-CUP1*). By contrast, crossover interference in the *msh4-6D* strain was similar to wild type, with the exception of one interval pair in which interference was diminished (*LYS5-MET13-CYH2*; **Figure 2F**). These exceptions suggest that the *msh4-6A* strain may not be completely defective for the formation of interfering crossovers, while *msh4-6D* may not be fully competent for this function.

Interference within individual intervals was also analyzed by calculating non-parental ditype (NPD) tetrad ratios (**Figure 2G** and **Supplemental Table S4**). Within a given interval, a double crossovers event involving all four chromatids results in a NPD tetrad. The NPD ratio compares the number of NPDs observed to that expected if there were no crossover interference (Papazian, 1952). A ratio of significantly less than one indicates positive crossover interference. Residual crossovers in the *msh4*Δ and *msh4-6A* strains did not show significant interference by NPD analysis. In the *msh4-6D* strain, interference was detected in all but one interval (*CEN3-MAT*). However, in wild-type tetrads, significant interference was detected in the same interval, again suggesting that *msh4-6D* is slightly defective for crossover interference

Next, we determined whether crossover assurance is influenced by Msh4 phosphorylation using a strain carrying eight linked intervals that span the length of chromosome III (**Figure 2H**)(Zakharyevich et al., 2010). In wild type, at least one crossover was detected in 98.9% of tetrads indicating highly efficient crossover assurance (**Figure 2I**). Oppositely, crossover assurance was severely defective in the absence of Msh4, with 25.2% of *msh4*Δ tetrads lacking a detectable crossover on chromosome III, consistent with previous analysis (Krishnaprasad et al., 2015). In addition, the fraction of tetrads with a single crossover was increased and multiple crossover classes were diminished in the *msh4*Δ strain relative to wild type (*P*<0.001, *G*-test). If crossover assurance remained operational in *msh4-6A* cells, the residual crossover frequency along chromosome III (1.7 crossovers per meiosis) is in principle sufficient to ensure a crossover in every meiosis. Contrary to this scenario, 17.4% of *msh4-6A* tetrads had zero crossovers highlighting the importance of Msh4 phosphorylation for crossover assurance (**Figure 2I**; *P*<0.001 compared to wild type, *G*-test; distributions of crossover classes were not different for *msh4-6A* and *msh4*Δ, *P*=0.38). The phospho-mimetic *msh4-6D* strain was not significantly different from wild type for crossover assurance, with just 2.8% of tetrads lacking a crossover (*P*=0.79). In conclusion, phosphorylation of the N-terminus of Msh4 promotes the formation of crossovers that are subject to patterning processes that result in crossover assurance and interference.

The contributions of individual phosphorylation sites to the crossover function of Msh4 were also assessed. This analysis revealed a major role for phosphorylation at sites S2, S4 and S7, while S41, T43 and S46 made little or no contribution to crossing over (**Supplemental Figure S3**). Western analysis indicated that Msh4-6A protein could still be phosphorylated, albeit with a delay and at lower levels than wild-type Msh4 (**Supplementary Figure S5;** low-level phosphorylation was also detected for Msh4-6D; also see **Figure 5A**;). This residual phosphorylation was abolished following mutation of all 18 serine and threonine residues present in the first 50 amino acids of Msh4 indicating that phosphorylation leading to the slow migrating form is confined to this region (**Supplementary Figure S5**). Importantly, the *msh4-18A* strain was no more defective for crossing over than the *msh4-6A* strain, indicating that phosphorylation of other sites in the N-terminus is not functionally redundant with the phosphorylation sites mapped by MS/MS (**Supplementary Figure S3**).

### Msh4 Phosphorylation Facilitates the Formation and Resolution of DNA Joint Molecules

To understand how the molecular steps of meiotic recombination are influenced by Msh4 phosphorylation, DNA intermediates were monitored in cultures undergoing synchronous meiosis using a series of Southern blot assays at the well-characterized *HIS4::LEU2* recombination hotspot (Hunter and Kleckner, 2001; Oh et al., 2007). At this locus, *Xho*I polymorphisms between the two parental chromosomes produce DNA fragments diagnostic for DSBs, JMs, and crossover products. DSBs, and crossovers were analyzed using one-dimensional (1D) gels (**Figure 3A–C**). Noncrossover products were detected by monitoring conversion of a *Bam*HI/*Ngo*MIV restriction-site polymorphism located directly at the site of DSB formation (**Figure 3A** and **3D**).

**Figure 3.**
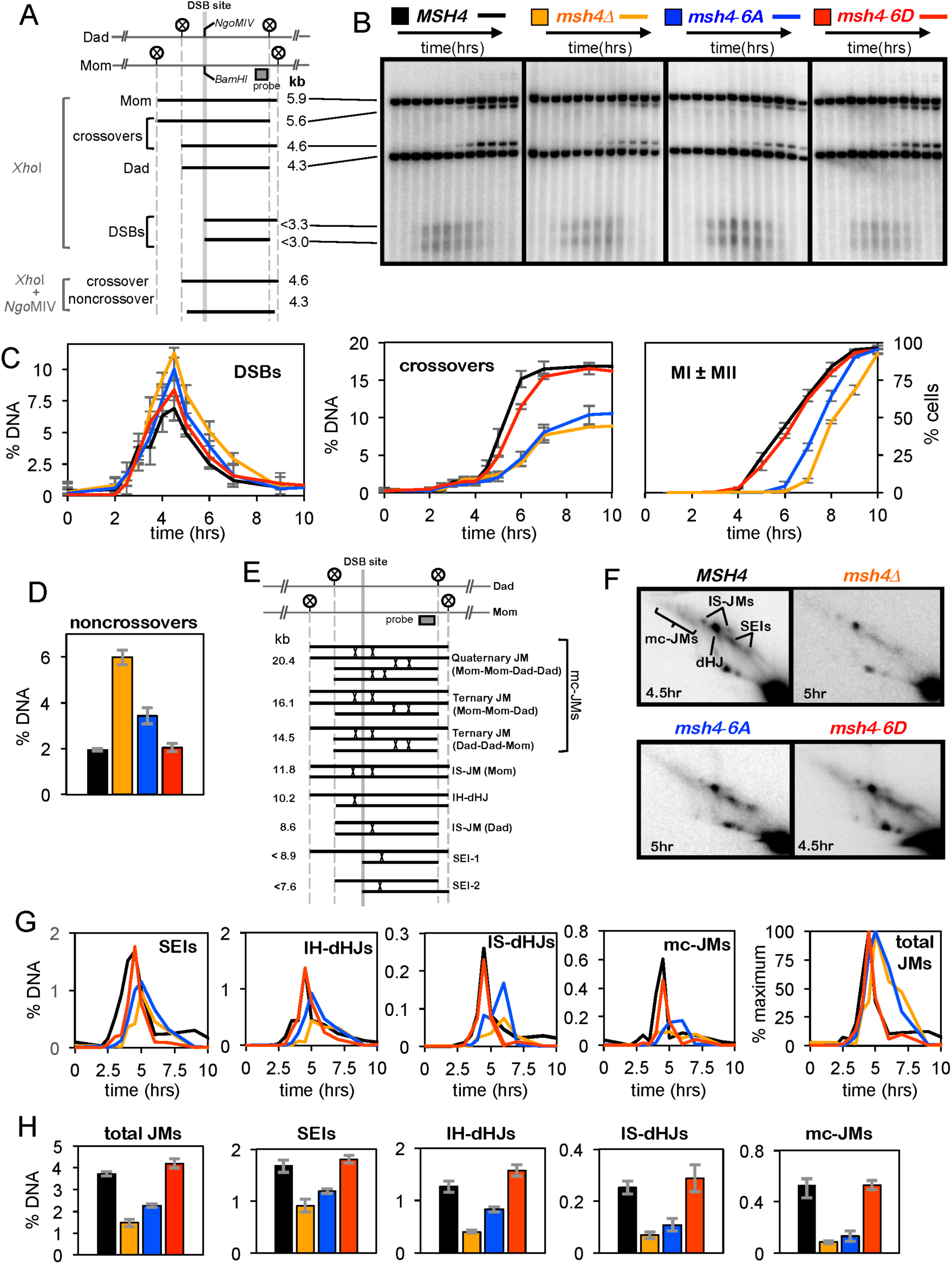
Physical analysis of the DNA events of meiotic recombination. (A) Map of *HIS4:LEU2* locus highlighting the DSB site, *Xho*I restriction sites (circled Xs) and the position of the probe used in Southern blotting. Sizes of diagnostic fragments are shown below. (B) Representative 1D gel Southern blot images for analysis of DSBs and crossovers. Time points are 0, 2, 2.5, 3, 3.5, 4, 4.5, 5, 6, 7, 9 and 11 hours. (C) Quantification of DSBs, crossovers and meiotic divisions. %DNA is percentage of total hybridizing DNA signal. MI ± MII is the percentage of cells that have completed one or both meiotic divisions. (D) Non-crossover levels at 11 hrs. (E) JM structures detected at the *HIS4:LEU2* locus. Positions of the DSB site, diagnostic *Xho*I sites (circled Xs) and the Southern probe are shown. (F) Representative 2D gel Southern blot images for time points where JM levels peak. Positions of the various JM signals are indicated in the first panel. (G) Quantification of JM species over time. (H) Quantification of JM species at their peak levels from three independent time courses. IH-dHJs, inter-homolog dHJs; IS-JMs, intersister JMs (most likely dHJs); SEIs, single-end invasions; mc-JMs, multi-chromatid JMs) and total JMs are indicated. Averages ± S.E. were calculated from three independent experiments.

Analysis of JM intermediates in budding yeast together with fine-scale analysis of recombination products in a variety of species, indicate that crossover and noncrossover pathways diverge at an early step, following nascent D-loop formation (Allers and Lichten, 2001b; Borner et al., 2004; Drouaud et al., 2013; Guillon et al., 2005; Hunter and Kleckner, 2001; Jeffreys and May, 2004; Marsolier-Kergoat et al., 2018; Martini et al., 2011; Rockmill et al., 2013; Wijnker et al., 2013). A majority of noncrossovers arise from D-loops via synthesis-dependent strand annealing in which the invading 3’ end is extended by DNA polymerase, unwound and then annealed to the other DSB end (Marsolier-Kergoat et al., 2018; McMahill et al., 2007). By contrast, most crossovers form via metastable one-ended strand-exchange intermediates called single-end invasions (SEIs), which form as homologs synapse (Hunter and Kleckner, 2001). Through DNA synthesis and second-end capture, SEIs give rise to double-Holliday junctions (dHJs)(Lao et al., 2008), which must then undergo biased resolution into crossovers (Zakharyevich et al., 2012). Native/native two-dimensional (2D) gels reveal the branched structure of JMs and were used to quantify SEIs, inter-homolog dHJs (IH-dHJs), inter-sister JMs (IS-JMs) and multi-chromatid JMs (mc-JMs) comprising three and four interconnected DNA molecules (**Figure 3E** and **3F**). To monitor the timing and efficiency of meiotic divisions, fixed cells were stained with DAPI and scored as having one, two, or four nuclei.

In wild-type, *msh4*Δ, *msh4-6A* and *msh4-6D* strains, DSBs appeared and reached peak levels with similar timing (**Figure 3C**). Peak DSB levels were higher and DSBs disappeared with a ∼1 hr delay in the *msh4*Δ null mutant relative to wild type, consistent with delayed progression of recombination and continued DSB formation (Borner et al., 2004; Thacker et al., 2014). Delayed progression in *msh4*Δ cells was also reflected by a ∼2 hr delay of the meiosis-I division (MI) (**Figure 3C**). Mirroring the crossover reductions detected by tetrad analysis, crossovers at *HIS4::LEU2* were reduced ∼2-fold in *msh4*Δ cells. A similar reduction in crossing over was seen for *msh4-6A* cells, but progression defects were less severe than those seen for *msh4*Δ; disappearance of DSBs was delayed by ∼30 minutes and MI was delayed by ∼1.25 hrs. In *msh4-6D* cells, slight delays (≤20 minutes) in DSB turnover and crossover formation were apparent, but crossovers reached wild-type levels.

The increased frequencies of gene conversion seen in *msh4*Δ and *msh4-6A* tetrads (**Figure 2E**) were mirrored by elevated levels of non-crossover gene conversions at *HIS4::LEU2* (**Figure 3D**). Again, the effect of *msh4-6A* was weaker than that of the *msh4*Δ null (increases of 1.8-fold versus 3.0-fold, respectively). Although gene conversion was also significantly elevated in *msh4-6D* tetrads, non-crossovers at *HIS4::LEU2* were not significantly increased.

Two-dimensional gel analysis revealed the importance of Msh4 phosphorylation for JM metabolism (**Figure 3E–H**). In *msh4-6A* cells, appearance of all JM species was delayed relative to wild-type by ∼30-60 min (**Figure 3G**), with the longest delays seen for JMs involving strand-exchange between sister chromatids, i.e. IS-JMs and mc-JMs. A further delay of ∼1.5 hrs was seen for the disappearance of JMs in *msh4-6A* relative to wild type. Peak JM levels were also lower in *msh4-6A* cells, averaging 61% of wild-type levels (**Figure 3H**). JM kinetics in *msh4*Δ null mutants were similar to those of *msh4-6A* (**Figure 3G**), but peak JM levels were significantly lower averaging just 40% of wild-type levels (**Figure 3H**). Thus, with respect to DSB persistence, JM levels and prophase delay, the phenotypes of the *msh4-6A* mutant are milder than those of the *msh4*Δ null, but still severely defective relative to wild type or *msh46D* strains.

Notably, SEIs reached similar levels in *msh4-6A* and *msh4*Δ cells (% of hybridizing DNA = 1.19% ± 0.05 S.E. and 0.91% ± 0.13 S.E., respectively), but dHJ levels were ∼2-fold lower in *msh4*Δ cells (0.83% ± 0.05 S.E. and 0.40% ± 0.02 S.E., respectively). Two non-exclusive possibilities could explain this difference: (i) the SEI-to-dHJ transition is less efficient and/or (ii) the stability of IH-dHJs is lower in the absence of Msh4 than when the Msh4-6A protein is present. However, despite higher IH-dHJ levels in *msh4-6A* cells, final crossover levels were very similar to those of the *msh4*Δ null (**Figures 2D, 2H** and **3C**). Together, these data suggest that phosphorylation of Msh4 is important both for JM formation and the crossover-biased resolution of IH-dHJs.

In *msh4-6D* cells, a minor delay in SEI formation was observed and IH-dHJs peaked at ∼24% higher levels relative to wild type (1.58% ± 0.11 versus 1.27% ± 0.11). But overall, JM kinetics and levels in *msh4-6D* cells were similar to those of wild type (**Figure 3G and 3H**).

### Msh4 Phosphorylation Facilitates Homolog Synapsis

In most organisms, the strand-exchange step of meiotic recombination promotes the pairing of homologs and their intimate end-to-end connection by synaptonemal complexes (SCs), meiosis-specific structures comprising densely-packed transverse filaments (Zickler and Kleckner, 2015). Crossovers then mature in the context of SCs, which are subsequently disassembled leaving homologs connected only at the sites of exchange. The possibility that the delayed and inefficient JM formation seen in *msh4-6A* cells leads to defective homolog synapsis was addressed by immuno-staining surface-spread nuclei for the synaptonemal complex transverse-filament protein, Zip1 (**Figure 4A** and **4B**)(Dong and Roeder, 2000). Synapsis was monitored over time by assigning nuclei to one of three classes based on the pattern of Zip1 staining (Borner et al., 2004): class I nuclei were defined by a dotty pattern; class II nuclei had partial synapsis with both linear and dotty staining; and class III had full synapsis indicated by extensive linear staining. Nuclei containing aggregates of Zip1 called polycomplexes (PCs), a sensitive indicator of synapsis defects (Sym and Roeder, 1995), were also quantified irrespective of their staining class.

**Figure 4.**
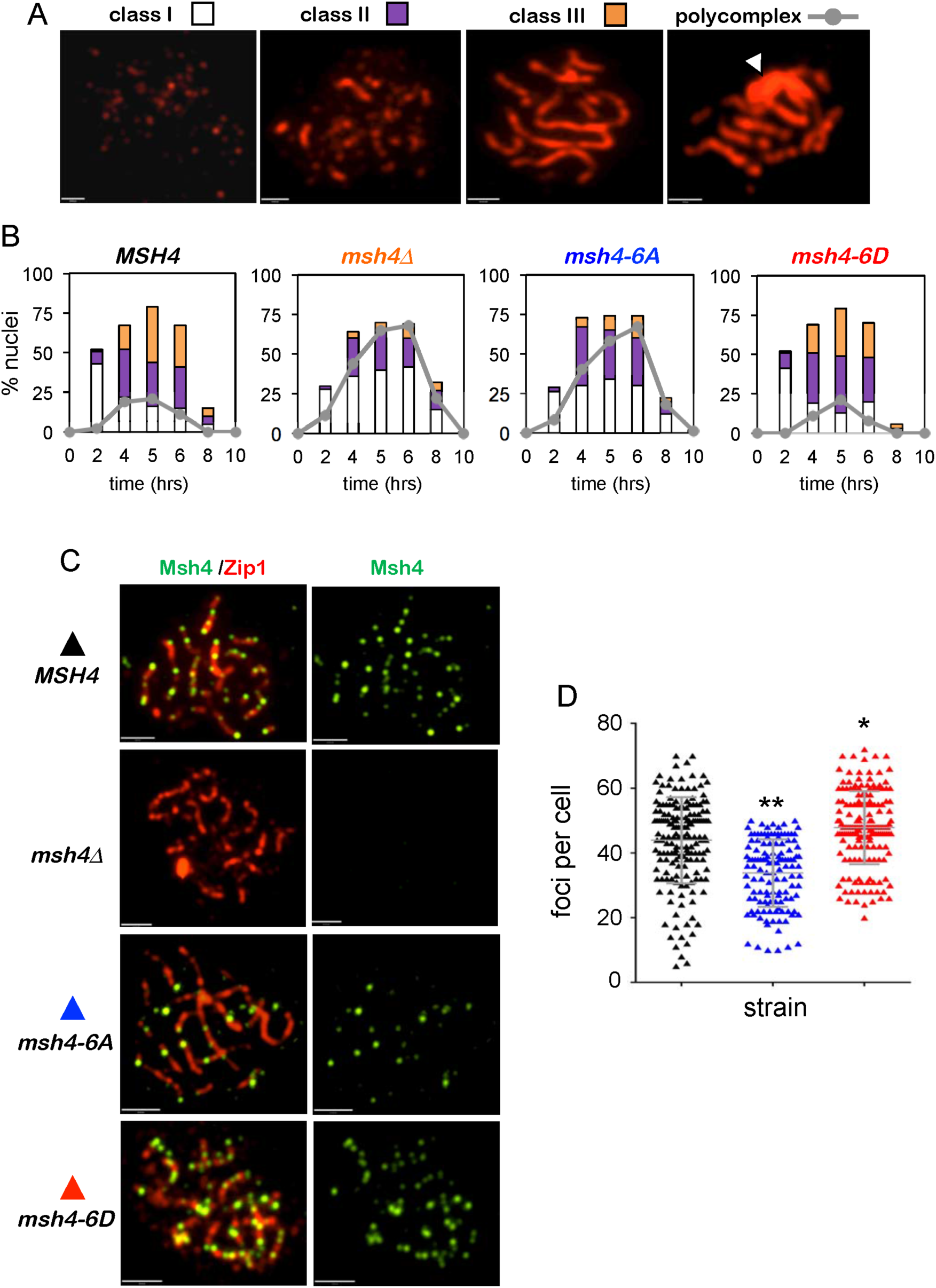
Chromosome synapsis and localization of Msh4 are facilitated by Msh4 phosphorylation. (A) Chromosome spreads showing representative examples of the three different Zip1 immuno-staining classes and a Zip1 polycomplex. (B) Quantification of Zip1 staining classes and polycomplexes. ≥100 nuclei were scored for each time point. (C) Representative images of spread meiotic nuclei immuno-stained for Msh4 (green) and Zip1 (red). (D) Quantification of Msh4 immuno-staining foci in class II and class III nuclei. ≥100 nuclei were scored for each strain. \**P*<0.05; \*\**P*<0.01 two tailed Mann Whitney test. Scale bars = 30 µm. Also see **Figure S4**.

In wild-type cells, peak levels of class III nuclei (35%) with full synapsis were seen at 5 hrs, and Zip1 has disappeared by 8 hrs (**Figure 4B**). Consistent with previous studies (Borner et al., 2004; Novak et al., 2001), synapsis was severely defective in *msh4*Δ null cells, with only 9% of cells achieving full synapsis and PCs present in a majority of cells (**Figure 4B**). PCs were similarly prominent in *msh4-6A* cells, but synapsis was slightly more efficient, with significantly higher levels of class II and class III nuclei (*P*<0.005, *G*-test). By contrast, synapsis in *msh4-6D* cells was indistinguishable from wild-type cells (*P* =0.63). Thus, Msh4 phosphorylation facilitates the formation and/or stabilization of SCs.

### Phosphorylation Promotes Chromosomal Localization of Msh4

To begin to understand how phosphorylation facilitates Msh4 function, the chromosomal localization patterns of Msh4, Msh4-6A and Msh4-6D proteins were compared. Surface spread nuclei were immunostained for both Msh4 and Zip1 (**Figure 4B** and **4C**). Msh4 foci were quantified in nuclei with zygotene (class II) and pachytene (class III) morphologies, i.e. partial and complete lines of Zip1 staining. In wild-type, Msh4 foci averaged 43.9 ± 13.3 S.D. per nucleus while focus numbers in *msh4-6A* nuclei were lower, averaging 33.8 ± 10.4 S.D. (*P*<0.0001, two-tailed Mann Whitney test; **Figure 4C**). By contrast, the Msh4-6D protein formed slightly elevated numbers of foci relative to wild-type Msh4, averaging 47.7 ± 11.3 S.D. per nucleus (*P*=0.028). Phenotypes associated with phosphorylation-defective (*msh4-3A*) and phosphorylation-mimetic (*msh4-3D*) alleles for sites S2, S4 and S7 were analogous to those of *msh4-6A* and *msh4-6D* with respect to formation of Msh4 foci (**Supplementary Figure S4**), further highlighting the importance of these three proximal serine residues.

### Msh4 is Stabilized by Phosphorylation

We explored the possibility that aberrant localization of phosphorylation-defective Msh4-6A protein is caused by decreased protein stability. Consistent with this idea, Western analysis showed that Msh4-6A protein levels were lower at all time points during meiosis, averaging a 2.2-fold reduction relative to wild-type Msh4 (**Figure 5A,B** and **Supplemental Figure S5**). In contrast, the Msh4-6D protein was hyper-stable, with an average increase of 2.1-fold. However, despite differences in steady-state protein levels, the overall timing of Msh4 appearance and disappearance was quite similar for Msh4, Msh4-6A and Msh4-6D proteins.

**Figure 5.**
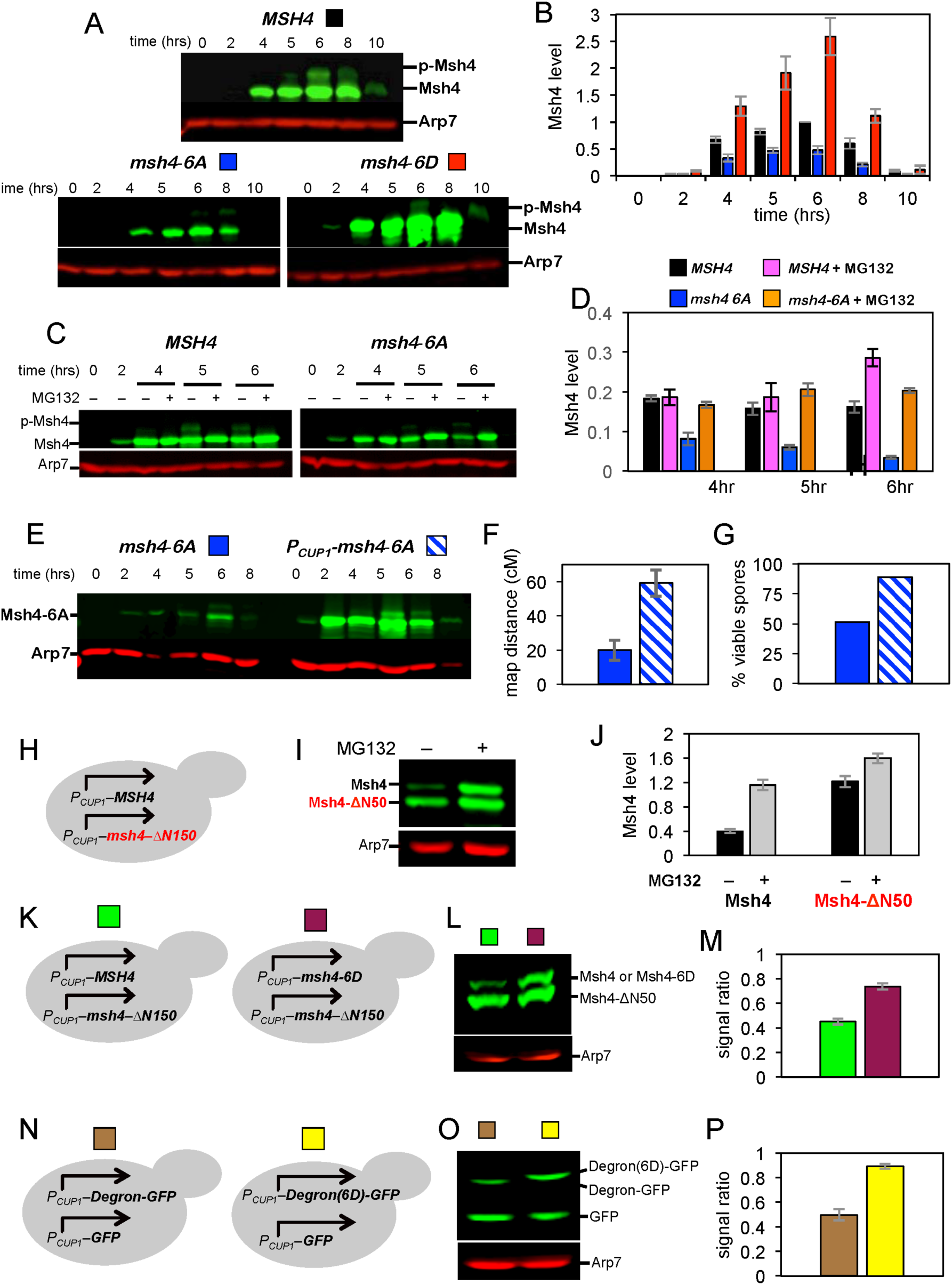
Degron activity of the Msh4 N-terminal region is attenuated by phosphorylation. (A) Western analysis of Msh4 during meiosis in wild-type, *msh-6A* and *msh4-6D* strains. (B) Quantification of Msh4 protein relative to the Arp7 loading control. Averages ± S.E. were calculated from three independent experiments. (C) Western analysis of Msh4 with and without addition of the proteasome inhibitor, MG132, at 2 hrs. (D) Quantification of Msh4 protein with and without MG132 treatment. Averages ± S.E. calculated from three independent experiments. (E) Western analysis of Msh4-6A protein during meiosis in the *msh4-6A* strain and following copper-induced overexpression in a *P*_*CUP1*_*-msh4-6A* strain. (F) Spore viability of *msh4-6A* and *P*_*CUP1*_*-msh4-6A* strains. (G) Map distances (± S.E.) for intervals flanking the *HIS4::LEU2* recombination hotspot (see **Figure S3**) in tetrads from *msh4-6A* and *P*_*CUP1*_*-msh4-6A* strains. (H) Experimental system for copper-inducible expression of Msh4 and msh4-ΔN50 proteins in vegetative cells. (I) Western analysis of the stains shown in panel E following copper induction, with and without MG132 treatment. (J) Quantification of the experiments represented in panels D and E. Averages ± S.E. were calculated from four independent experiments. (K) Experimental systems to compare co-expression of Msh4 and Msh4-δN50 (left), with co-expression of Msh4-6D and Msh4-δN50 proteins (right). (L) Western analysis of the stains shown in panel H following copper induction, with and without MG132 treatment. (M) Quantification of experiments represented in panels H and I. Average ratios ± S.E. were calculated from four independent experiments. (N) Experimental systems for co-expression of GFP and the Msh4 N-terminal region (“degron”) fused to GFP (left); or co-expresson of GFP and a phospho-mimetic derivative of the Msh4 N-terminal region (“degron 6D) fused to GFP. (O) Western analysis of the stains shown in panel K following copper induction. (P) Quantification of experiments represented in panels K and L. Average ratios ± S.E. were calculated from four independent experiments. Also see **Figure S5**.

Next, we examined whether the lower level of the Msh4-6A protein was due to proteasome-mediated degradation. Wild-type and *msh4-6A* cells were treated with the proteasome inhibitor MG132 two hours after transfer to sporulation medium and Msh4 protein levels were measured at 4, 5 and 6 hrs by Western blot (**Figure 5C,D**). In the absence of MG132, Msh4-6A protein levels were reduced to 21-44% of wild-type levels. Treatment with MG132 restored Msh4-6A levels to between 94% and 125% of Msh4 levels seen in control wild-type cells. MG132 treatment did not have a significant effect on wild-type Msh4 levels at 4 and 5 hrs, but at 6 hrs levels were 1.8-fold higher than in untreated cells (**Figure 5C,D**). These data imply that phosphorylation stabilizes Msh4 during meiotic prophase I by protecting it from proteasomal degradation.

If the primary function of phosphorylation is to stabilize Msh4, then overexpression of phosphorylation-defective Msh4-6A protein should suppress *msh4-6A* mutant phenotypes. Indeed, overexpression of *msh4-6A* using the strong, copper-inducible *CUP1* promoter restored crossing-over and spore viability to near wild-type levels (**Figure 5E–G**).

### The Msh4 N-Terminus Encodes a Portable Degron

Our analysis points to a model in which Msh4 is an intrinsically unstable protein that is stabilized by phosphorylation of N-terminal residues, thereby activating the crossover function of MutSγ. To further test this model, full-length Msh4 and an N-terminally truncated derivative (Msh4-ΔN50) were co-expressed in vegetative (non-meiotic) yeast cells using the *CUP1* promoter (**Figure 5H–J** and **Supplemental Figure S5**). The steady-state level of full-length Msh4 was 3-fold lower than that of Msh4-ΔN50 and treatment with MG132 showed that this difference was due to proteasomal degradation. By contrast, the N-terminus of Msh5 had no effect on its stability (**Supplemental Figure S6**). Thus, the N-terminal region of Msh4 possesses degron activity.

Comparison of protein levels in strains co-expressing wild-type Msh4 and Msh4-ΔN50 versus Msh4-6D and Msh4-ΔN50 revealed that the phospho-mimetic allele significantly attenuated N-terminal degron activity (**Figure 5K–M**). Protein half-lives, estimated from cycloheximide chase experiments, were ∼14, 31 and 61 minutes respectively for Msh4, Msh4-6D and Msh4-ΔN50 (**Supplemental Figure S5**).

To address whether the Msh4 N-terminus has autonomous, portable degron activity, residues 1-50 of wild-type Msh4 (“Degron”) or a phospho-mimetic derivative (“Degron(6D)”) were fused to GFP and co-expressed in vegetative cells together with wild-type GFP (**Figure 5N**). The Msh4degron destabilized GFP, reducing its half-life from ∼59 to 19 mins (**Figure 5O,P** and **Supplemental Figure S5**). By contrast, stability of the phospho-mimetic Degron(6D)-GFP fusion was similar to that of wild-type GFP (half life of ∼49 mins). Together, these data indicate that the N-terminal domain of Msh4 comprises an autonomous degron that is neutralized by phosphorylation.

### Msh4 Phosphorylation Occurs *In Situ* At Sites of Recombination

The genetic requirements for Msh4 phosphorylation were delineated by performing Western analysis on extracts from mutant strains that are defective for successive steps in meiotic recombination, chromosome pairing and synapsis (**Figure 6A**). The slow migrating band diagnostic of phosphorylated Msh4 was not detected in *spo11-Y135F*, *mnd1*Δ, *zip1*Δ and *zip3*Δ mutants (**Figure 6B**). The *spo11-Y135F* allele lacks the catalytic tyrosine required for DSB formation and therefore fails to initiate meiotic recombination (Bergerat et al., 1997). Mnd1 is an essential co-factor for DNA strand-exchange and *mnd1*Δ mutant cells are severely defective for DSB repair, homolog pairing and synapsis (Chen et al., 2004; Gerton and DeRisi, 2002; Tsubouchi and Roeder, 2002). *zip1*Δ cells achieve homolog pairing, but formation of crossover-designated JMs is defective and synapsis fails because Zip1 is the major component of the SC central region (Borner et al., 2004; Sym et al., 1993). The SUMO E3 ligase, Zip3, accumulates at future crossover sites and facilitates the loading of other ZMM factors, including MutSγ (Agarwal and Roeder, 2000; Cheng et al., 2006; Henderson and Keeney, 2004; Shinohara et al., 2008). *zip3*Δ mutants are defective for homolog synapsis, JM formation and crossing over (Agarwal and Roeder, 2000; Borner et al., 2004). Thus, Msh4 phosphorylation is DSB-dependent and requires synapsis and the formation of crossover designated JMs.

**Figure 6.**
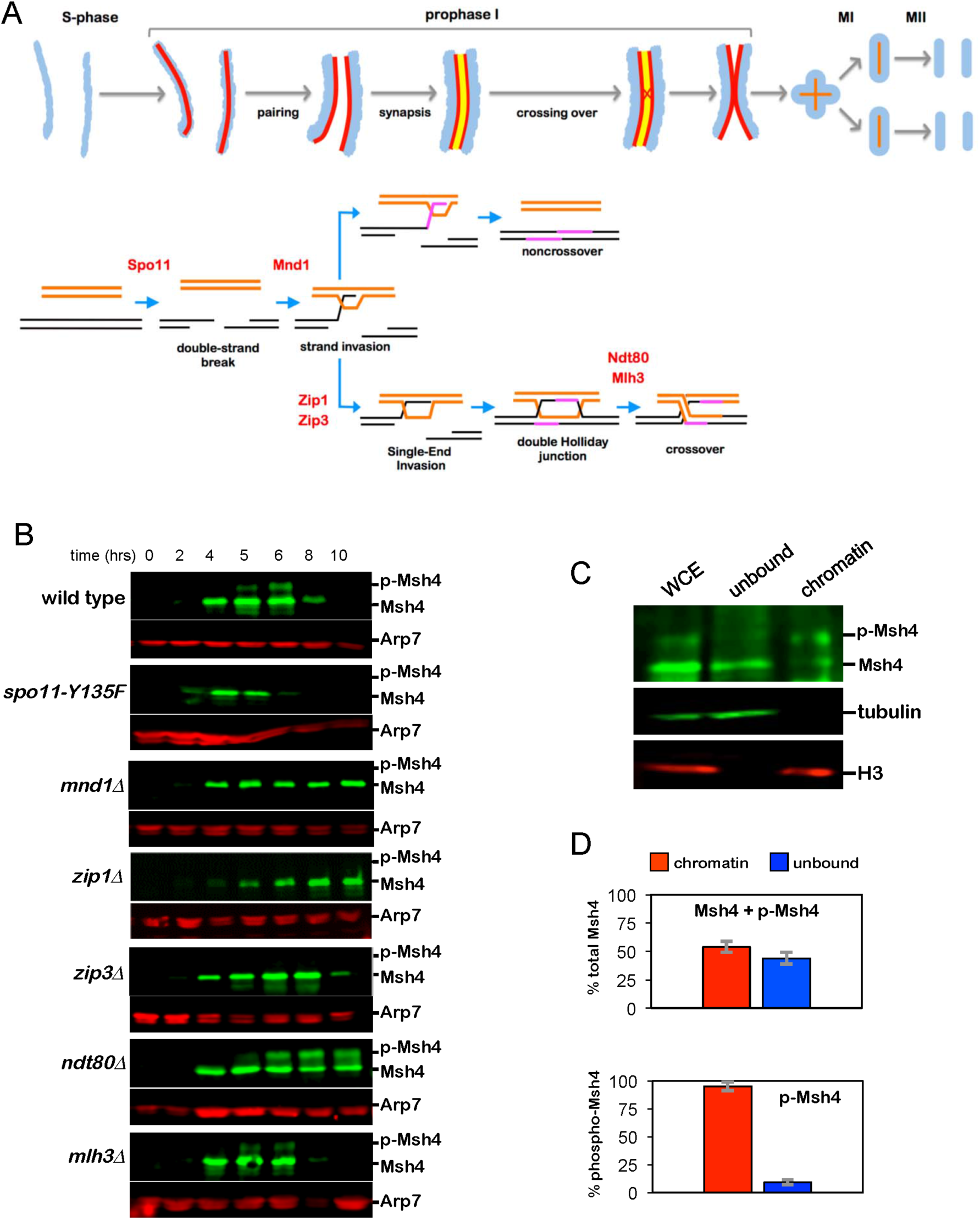
Genetic requirements and chromatin association of phosphorylated of Msh4. (A) Chromosomal and recombination events of meiosis illustrating the steps affected by mutants analyzed in panel B. Blue, chromatin; red lines, homolog axes; yellow line, synaptonemal complex central region. (B) Western analysis of Msh4 in the indicated mutants. (C) Western analysis of Msh4 from whole cell extracts (“WCE”) and extracts separated into soluble (“unbound”) and chromatin fractions; tubulin and histone H3 were used as markers for these two fractions, respectively. (D) Quantification of total Msh4 (top graph) and phosphorylated Msh4 (bottom graph) in the two fractions. Means values ±S.E. were calculated from three independent experiments.

Analysis of two additional mutants, *ndt80*Δ and *mlh3*Δ, showed that Msh4 phosphorylation does not require dHJ resolution or crossing over (**Figure 6B**). Ndt80 is a transcription factor required for meiosis to progress beyond pachytene (Chu and Herskowitz, 1998). *ndt80*Δ cells arrest with fully synapsed chromosomes and unresolved JMs (Allers and Lichten, 2001a; Xu et al., 1995). In *ndt80*Δ cells, phospho-Msh4 accumulated to high levels (≥60% of total Msh4) and persisted in arrested cells (**Figure 6B**). The MutLγ complex, comprising MutL homologs Mlh1 and Mlh3, possesses endonuclease activity that is required for the biased resolution of dHJs into crossovers, but not for JM resolution *per se* (Claeys Bouuaert and Keeney, 2017; Manhart et al., 2017; Ranjha et al., 2014; Rogacheva et al., 2014; Zakharyevich et al., 2012). In *mlh3*Δ mutants, chromosomes synapse and normal levels of JMs are formed and resolved, but crossing over is defective (Zakharyevich et al., 2012). Phosphorylation and turnover of Msh4 appeared normal in *mlh3*Δ cells (**Figure 6B**).

Overall, our analysis suggests that Msh4 is phosphorylated *in situ* at sites of recombination, likely when bound to JM intermediates in the context of synapsed chromosomes. Consistent with this inference, chromatin-associated Msh4 was highly enriched for the phosphorylated form relative to soluble Msh4 (**Figure 6C,D**). Specifically, while total Msh4 protein was roughly equally distributed between chromatin-bound and soluble cell fractions, 95% of phospho-Msh4 was found in the chromatin-bound fraction (**Figure 6C,D**).

### Msh4 Phosphorylation is Catalyzed by Dbf4-Dependent Kinase, Cdc7

To identify the kinase(s) responsible for Msh4 phosphorylation, the requirement for candidate kinases was systematically analyzed (**Figure 7**). The PI3K-like kinases, Mec1^ATR^ and Tel1^ATM^ are primary sensor kinases of the DNA-damage response and function in meiosis to regulate DSB distribution, inter-homolog template bias and crossing over (Carballo and Cha, 2007; Cooper et al., 2016). Msh4 phosphorylation was reduced more than 2-fold in *P*_*CLB2*_*-MEC1* cells in which the essential *MEC1* gene is expressed under the meiotically-repressed *CLB2* promoter (**Figure 7A** and **B**)(Lee and Amon, 2003). In contrast, a kinase dead mutant of *TEL1* had no effect on Msh4 phosphorylation (**Figure 7A**, (Ma and Greider, 2009)). However, combining *P*_*CLB2*_*-MEC1* with *tel1-kd* abolished Msh4 phosphorylation and lowered protein levels of Msh4 indicating that *TEL1* can partially compensate for the lack of *MEC1* with regard to Msh4 phosphorylation.

**Figure 7.**
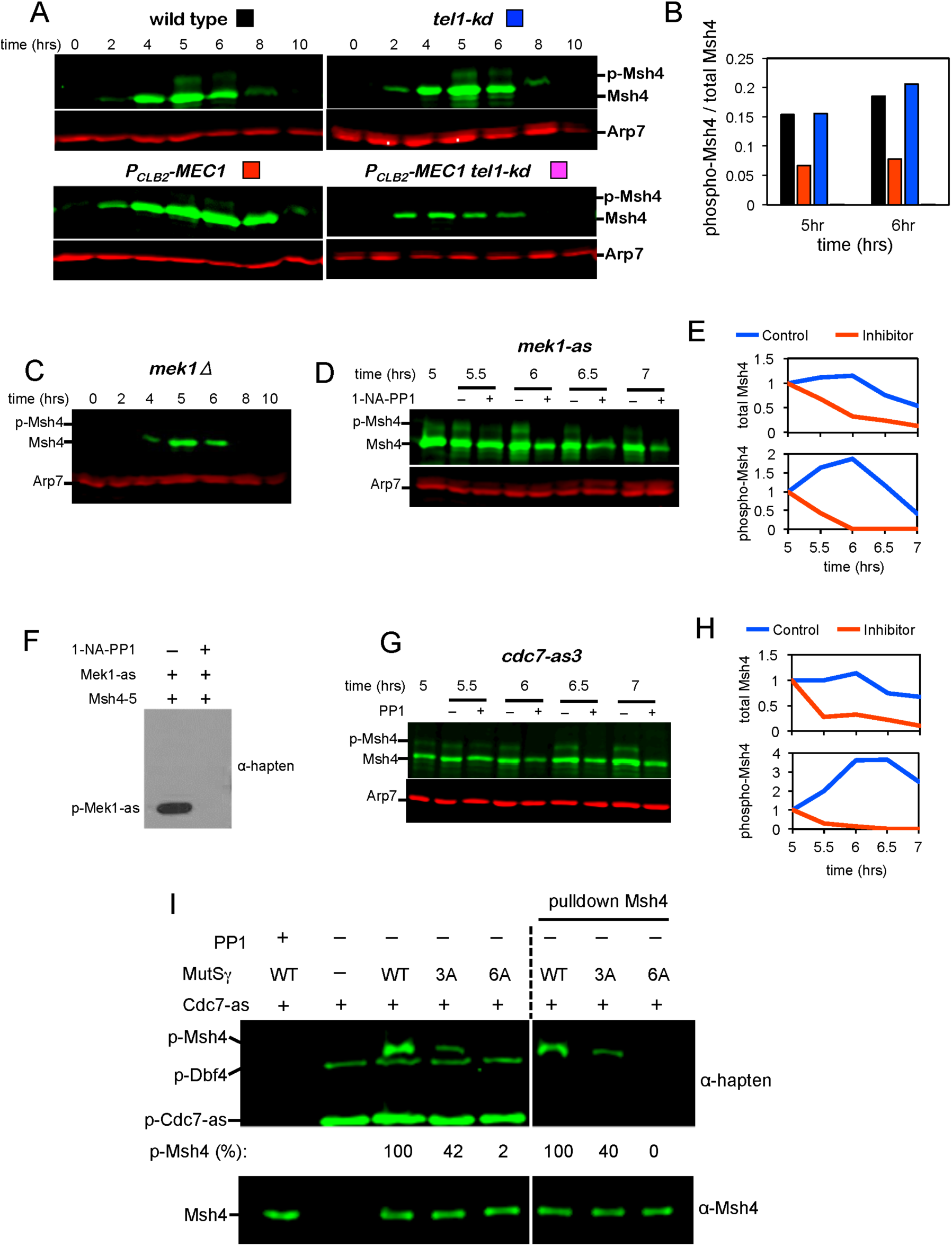
Kinase requirements for phosphorylation of Msh4 *in vivo* and *in vitro.* (A) Western analysis of Msh4 in *tel1-kd, pCLB2-MEC1* and *pCLB2-MEC1 tel1-kd* strains during meiosis. (B) Fraction of Msh4 that is phosphorylated at 5 and 6 hrs in the experiments shown in panel (A). (C) Western analysis of Msh4 in a *mek1*Δ null mutant. (D) Western analysis of Msh4 in strains containing the ATP-analog sensitive *mek1-as* allele, with and without addition of the inhibitor 1-NA-PP1 at 5 hrs. (E) Relative levels of total and phosphorylated Msh4 quantified from the experiment shown in panel (D). Levels were normalized to the 5 hr time point. (F) Western analysis of a Mek1-as (GST-Mek1-as) *in vitro* kinase assay with MutSγ, with and without the inhibitor 1-NA-PP1. (G) Western analysis of Msh4 in strains containing the ATP-analog sensitive *cdc7-as* allele, with and without addition of the inhibitor PP1 at 5 hrs. The anti-thiophosphate ester (α-hapten) antibody recognizes phosphorylation products of the semi-synthetic epitope system. (H) Relative levels of total and phosphorylated Msh4 quantified from the experiment shown in panel (G). Levels were normalized to the 5 hrs timepoint. (I) Western analysis of an *in vitro* kinase assay with the Cdc7-as3 kinase (FLAG-Cdc7-as3– Dbf4 complex) with wild-type MutSγ or mutant derivatives containing serine-alanine substitutions in the N-terminus of Msh4. In the right-hand panel, Msh4 was purified from the reactions. Phosphorylation efficiency was calculated relative to wild-type Msh4. In the lower panels, 10% of each reaction was probed for Msh4 as a loading control. Also see **Figure S6**.

None of the identified Msh4 phosphorylation sites conform to the S/T-Q Mec1/Tel1 target-site consensus (**Figure 1E**), making it unlikely to be a direct target. Thus, we explored whether signaling pathways downstream of Mec1/Tel1 are important for Msh4 phosphorylation. An important meiotic target of Mec1/Tel1 is Hop1, a checkpoint adaptor for the meiotic DNA damage response that recruits and activates the serine/threonine effector kinase, Mek1 (Carballo et al., 2008; Niu et al., 2005). Mek1 promotes inter-homolog bias, synapsis and crossing over, and regulates meiotic progression (Callender et al., 2016; Chen et al., 2015; Hollingsworth, 2010; Niu et al., 2007; Prugar et al., 2017; Subramanian et al., 2016; Wu et al., 2010). Although Mek1 consensus-target sites (RXX<u>T</u>) are absent from the Msh4 N-terminus, a *mek1*Δ null mutation abolished Msh4 phosphorylation and reduced steady-state protein levels (**Figure 7C**). Diminished Msh4 phosphorylation in *mek1*Δ cells could be an indirect effect of the defective inter-homolog interactions caused by this mutation. Therefore, we employed a chemical genetic approach to inhibit Mek1 kinase activity after inter-homolog interactions and synapsis had been established (**Figure 7D**). Inhibition of the analog-sensitive *mek1-as* allele (Wan et al., 2004) at 5 hrs after transfer to sporulation medium resulted in the rapid disappearance of phosphorylated Msh4 and a reduction in Msh4 protein levels (**Figure 7D** and **E**). To test whether Msh4 is a direct substrate of Mek1, recombinant MutSγ was incubated with immuno-purified GST-Mek1-as (**Figure 7F**). Phosphorylation catalyzed specifically by GST-Mek1-as was detected using the ATPγS analog, N^6^-Furfuryl-ATPγS, and the semi-synthetic epitope system (Allen et al., 2007; Lo and Hollingsworth, 2011). Under these conditions, only auto-phosphorylation of Mek1-as was detected indicating that neither Msh4 or Msh5 is a direct target of Mek1.

Dbf4-dependent kinase (DDK) comprises the kinase Cdc7 and regulatory subunit Dbf4 (Matsumoto and Masai, 2013). DDK functions throughout meiosis and promotes ZMM-mediated crossing over through phosphorylation of Zip1 (Chen et al., 2015). DDK prefers to phosphorylate serines and threonines immediately upstream of a negative charge, which can be conferred either by negatively charged amino acids, such as aspartate or glutamate, or by phosphorylation (Cho et al., 2006; Montagnoli et al., 2006). Msh4 S2 and S7 and S46 are candidates for DDK sites bases on these criteria (**Figure 1E**). Analogous to *mek1-as,* inhibition of the analog-sensitive *cdc7-as* allele (Wan et al., 2006) at 5 hrs (after Cdc7 has activated DSB and SC formation) caused rapid disappearance of phosphorylated Msh4 accompanied by a reduction in total protein level (**Figure 7G** and **H**).

*In vitro* phosphorylation reactions showed that DDK directly phosphorylates the Msh4 N-terminus (**Figure 7I**). To specifically detect phosphorylation catalyzed by Cdc7, immuno-purified FLAG-Cdc7-as/Dbf4 complex was incubated with 6-Benzyl-ATPγS and phosphorylation was detected via the semi-synthetic epitope system (Allen et al., 2007; Chen et al., 2015; Lo and Hollingsworth, 2011). In the absence of recombinant MutSγ, only auto-phosphorylation of Cdc7-as and Dbf4 was detected. When wild-type MutSγ (“WT” in **Figure 7I**) was added, an additional phosphorylation product was detected that could be either Msh4 or Msh5 based on its molecular weight. The identity of this product was determined in two ways. First, MutSγ derivatives containing Msh4-3A (“3A” mutant for S2, S4 and S7) or Msh4-6A (“6A”) proteins were used as substrates. Phosphorylation of Msh4-3A was reduced by 68% relative to wild type, while Msh4-6A was not modified by DDK, clearly identifying Msh4 as the DDK substrate (**Figure 7I**).

Second, when Msh4, Msh4-3A and Msh4-6A proteins were purified from DDK kinase assay mixtures, only the largest phosphorylation product was detected. In addition, all phosphorylation products were sensitive to inhibition of Cdc7-as by the bulky ATP analog, PP1, confirming the specificity of this reaction.

## DISCUSSION

### Regulated Proteolysis Is a Key Aspect of Meiotic Crossing Over

The molecular mechanisms that underpin the differentiation of meiotic crossover and non-crossover pathways have remained elusive. Specifically, it is not known how events leading to dHJ formation are facilitated at some recombination sites but not at others, and how dHJs maintain their crossover fate and undergo crossover-biased resolution. Here, regulated proteolysis is revealed as a key determinant of crossing over. This discovery substantiates previous studies implicating the ubiquitin-proteasome system (UPS) in crossover/non-crossover differentiation (Ahuja et al., 2017; Qiao et al., 2014; Rao et al., 2017; Reynolds et al., 2013). Notably, when the UPS is inactivated in mouse spermatocytes, meiotic recombination stalls and ZMM factors (including MutSγ) persist at sites that would normally mature into non-crossovers, suggesting that ZMMs may be targeted for proteolysis at these sites (Rao et al., 2017). At least one ZMM factor, Msh4, can now be designated as a direct target of proteasomal degradation. The atypical mode of Msh4 regulation reveals unanticipated facets of crossover differentiation: intrinsic instability of an essential factor dictates that non-crossover will be the default outcome, and kinase-dependent stabilization activates crossing over.

### The Crossover Activity of MutSγ Is Activated by Stabilizing Msh4

Distinct activities of the ZMMs influence different aspects of crossover maturation and couple these events to homolog synapsis. The DNA helicase, Mer3, functions both to regulate the extension of nascent D-loops by DNA synthesis and stabilize JMs (Borner et al., 2004; Duroc et al., 2017; Mazina et al., 2004; Nakagawa and Kolodner, 2002); the XPF-ERCC1 related complex, Zip2-Spo16, specifically binds JMs (De Muyt et al., 2018; Guiraldelli et al., 2018; Macaisne et al., 2011); Zip1 acts both locally to promote ZMM function and globally as the major component of SCs (Chen et al., 2015; Sym et al., 1993; Voelkel-Meiman et al., 2015); Zip3 is a SUMO E3 ligase that helps localize other ZMMs to nascent crossover sites and facilitates synapsis (Agarwal and Roeder, 2000; Cheng et al., 2006; Macqueen and Roeder, 2009; Shinohara et al., 2008); and Zip4 is a large TPR repeat protein thought to bridge interactions between Zip2-Spo16, Zip3, MutSγ and the chromosome axis protein Red1 (De Muyt et al., 2018). Several activities are ascribed to MutSγ: (i) specific binding to JM structures (D-loops and Holliday junctions)(Snowden et al., 2004); (ii) stabilization of nascent JMs following ATP-dependent conversion of JM-bound MutSγ into sliding clamps that diffuse away from junction points while embracing two DNA duplexes (Snowden et al., 2004); (iii) protection of dHJs from the anti-crossover “dissolution” activity of the STR decatenase complex, Sgs1–Top3–Rmi1 (equivalent to the human BTR complex, BLM–TOPIIIα–RMI1/2)(Jessop et al., 2006; Kaur et al., 2015; Oh et al., 2007; Tang et al., 2015)(Tang and Hunter, unpublished); (iv) direct or indirect recruitment and activation of crossover-biased JM resolving factors such as the MutLγ endonuclease (Manhart et al., 2017; Nishant et al., 2008; Ranjha et al., 2014; Zakharyevich et al., 2012); (v) formation and/or stabilization of homolog synapsis (Borner et al., 2004; Novak et al., 2001).

Phosphorylation-defective Msh4-6A protein can still localize to chromosomes and retains significant function for synapsis and JM formation. However, the essential crossover function(s) of MutSγ is inactive unless Msh4 is stabilized via phosphorylation. We suggest that these essential functions are to protect dHJs from STR/BTR-mediated dissolution and facilitate their biased resolution. This proposal is also consonant with our inference that DDK targets MutSγ complexes that have bound JMs in the context of synapsed or synapsing chromosomes. Notably, STR/BTR complexes also accumulate at crossover sites (Jagut et al., 2016; Rockmill et al., 2003; Woglar and Villeneuve, 2018); and the symmetric arrangement of dual foci of MutSγ and BTR observed in *C. elegans* suggests a specific model in which MutSγ sliding clamps accumulate between the two junctions of a dHJ to impede dissolution (Woglar and Villeneuve, 2018). We propose that sliding clamps of MutSγ must accumulate above a minimum number to facilitate crossing over, be it through dHJ stabilization, recruitment or activation of resolving enzymes, or maintaining dHJs in a geometry that is conducive to crossover-biased resolution. Under this model, the requisite threshold of dHJ-bound MutSγ clamps requires the stabilization of Msh4 by phosphorylation. We note that the estimated half-life of stabilized Msh4 (30-60 mins) is similar to the estimated lifespan of dHJs (Allers and Lichten, 2001a; Hunter and Kleckner, 2001) suggesting a causal relationship.

If MutSγ were the primary limiting factor for crossing over, then the hyper-stable phospho-mimetic Msh4-6D protein would be expected to increase crossing over. Although *msh4-6D* strains showed modest increases in Msh4 foci and IH-dHJs, elevated gene conversion and mildly perturbed interference, crossing over was not increased. Thus, some other factor(s) limits crossover numbers. The most likely interpretation is that crossovers are limited by interference and stabilization of Msh4 occurs at designated crossover sites, downstream of the initial crossover/non-crossover decision. However, the proteolysis mechanism revealed here for Msh4 could be a general mechanism to regulate the availability of essential crossover factors and thereby limit crossovers.

We further suggest that the intrinsic instability of Msh4 may be enhanced by proximity to proteasomes, which are recruited in high numbers along chromosome axes as they synapse (Ahuja et al., 2017; Rao et al., 2017). This SC-associated population of proteasomes could accelerate the loss of MutSγ from synapsed sites where Msh4 is not stabilized by phosphorylation and thereby drive recombination towards a non-crossover outcome.

Whether MutSγ is similarly regulated in other organisms remains unclear. An N-terminal region appears to be common to all Msh4 proteins, but sequence conservation is low. However, these regions are typically S/T rich, contain candidate DDK sites and are predicted to undergo disorder-enhanced phosphorylation (**Supplemental Figure S1** and data not shown). The intrinsic stability of Msh4 would presumably have to coevolve with the duration of meiotic prophase in different species, which can vary by at least an order of magnitude.

### DDK Is A Key Effector of Meiotic Prophase

In addition to stabilizing Msh4 to activate MutSγ for crossing over, DDK facilitates meiotic S-phase (Valentin et al., 2006; Wan et al., 2006); triggers DSBs and couples their formation to the passage of replication forks (Matos et al., 2008; Murakami and Keeney, 2014; Sasanuma et al., 2008; Wan et al., 2008); promotes synapsis and crossing over via phosphorylation of Zip1 on its C-terminus (Chen et al., 2015); enables progression beyond pachytene by removing the Sum1 repression complex from the *NDT80* promoter (Lo et al., 2012; Lo et al., 2008); drives the destruction of SCs (Argunhan et al., 2017); is required to recruit monopolin to kinetochores enabling mono-orientation of homologs on the meiosis-I spindle (Lo et al., 2008; Matos et al., 2008); and facilitates the cleavage of cohesin to allow homolog disjunction at the meiosis-I division (Katis et al., 2010). Thus, DDK is a primary effector kinase for all the major events of meiotic prophase.

Direct targeting of both Zip1 and Msh4 implies that DDK is a general activator of ZMM-mediated crossing over. However, the timing, requirements and modes of regulation are distinct. By contrast to Msh4, Zip1 phosphorylation is an early event that depends on DSB formation but not later steps of recombination and doesn’t act by stabilizing the protein (Chen et al., 2015). Moreover, DDK-mediated phosphorylation of Zip1 is inferred to function upstream of the other ZMMs. Consistent with this inference, Msh4 phosphorylation requires both the presence of Zip1 and phosphorylation of its C-terminus (**Figure 6** and data not shown). Importantly, the upstream requirement for Mek1 in DDK-mediated Zip1 phosphorylation (Chen et al., 2015) explains why Msh4 phosphorylation is also Mek1 dependent.

It’s unclear how DDK achieves successive, dependent phosphorylation of Zip1 and Msh4. An intriguing possibility is that DDK is sequestered by Zip1 until synapsis ensues, a model suggested by recent analysis of substrate ordering in mitotically cycling cells (Seoane and Morgan, 2017). Alternatively, the N-terminal region of Msh4 could be masked until MutSγ converts to a sliding clamp on DNA; or crossover-designated recombination complexes could create composite docking sites for DDK. Apparent ordering could involve rapid reversal of DDK-catalyzed phosphorylation until Msh4 becomes protected at designated crossover sites. With this model in mind, Woglar and Villeneuve recently demonstrated that crossover-designated recombination complexes become enveloped in “bubbles” of SC central region proteins that could protect components from both phosphatases and proteasomes (Woglar and Villeneuve, 2018).

### Contingent Kinase Cascades Order The Events of Meiotic Prophase

While DKK appears to be the ultimate effector for many prophase events, other kinases dictate its activity in space and time. CDK primes DDK phosphorylation of Mer2 to trigger DSB formation (Henderson et al., 2006; Sasanuma et al., 2008; Wan et al., 2008). DSB-dependent activation of Mec1^ATR^/Tel1^ATM^ locally activates Mek1 (Carballo et al., 2008), which promotes inter-homolog recombination via its direct targets (Callender et al., 2016; Niu et al., 2009), and indirectly activates synapsis and the ZMM pathway by licensing DDK to phosphorylate Zip1 and subsequently Msh4 (Chen et al., 2015)(this study). CDK, DDK and the meiosis-specific kinase Ime2 collectively target the Sum1 transcriptional repressor to help activate expression of the transcription factor Ndt80 and exit from pachytene. The polo-like kinase Cdc5, whose expression is Ndt80 dependent, then collaborates with CDK and DDK to disassemble SCs (Argunhan et al., 2017; Sourirajan and Lichten, 2008). Cdc5 also collaborates with DDK to localize the monopolin complex to MI kinetochores (Lo et al., 2008; Matos et al., 2008). Finally, casein kinase δ/ε works with DDK to activate the cleavage of cohesin by separase and trigger the meiosis-I division (Katis et al., 2010). Understanding the spatial-temporal regulation of DDK with respect to the activation of ZMM-dependent crossing over, and its relationship to crossover control are important goals for the future.

## EXPERIMENTAL PROCEDURES

Extended methods are described in the Supplemental Information.

## SUPPLEMENTAL INFORMATION

Supplemental Information includes 7 figures, 5 tables and extended methods.

## ACKNOWLEDGEMENTS

We thank members of the Hunter Lab for support and discussions. This work was supported by NIH NIGMS grant GM074223 to N.H. and GM050717 to N.M.H. S.T. was supported by an NIH NIEHS-funded training program in Environmental Health Sciences (T32 ES007058). N.H. is an Investigator of the Howard Hughes Medical Institute.

## AUTHOR CONTRIBUTIONS

W.H. and N.H. conceived the study and designed most experiments. N.M.H., X.C. and W.H. designed *in vitro* kinase experiments. All authors performed experiments and analyzed the data. W.H. and N.H. wrote the manuscript with inputs and edits from all authors.

